# Structural Consequences of the Villin Headpiece Interaction with a Carbon Nitride Polyaniline (C_3_N) Nanosheet

**DOI:** 10.1101/2020.06.17.158402

**Authors:** Zonglin Gu, Jose Manuel Perez-Aguilar, Ruhong Zhou

## Abstract

Carbon nitride polyaniline (C_3_N) nanosheets shared a similar structure with graphene and have been utilized in biomedical applications since its recent successful synthesis. However, limited information was known about the interaction of this next-generation nanomaterial with biomolecules, which might hamper its applications in living tissues. Here, by using all-atom molecular dynamics (MD) simulations, we investigated the interaction between a C_3_N nanosheet and the prototypical protein villin headpiece (HP35), in order to identify the mechanistic determinants of such interaction; this knowledge will provide guidelines about C_3_N’s biocompatibility. Our MD simulations revealed that the C_3_N-based nanomaterial caused the partial denaturation of HP35 once the protein was bound on its surface. That is, upon adsorption, we observed the loss of the protein’s interior hydrogen bonds and the native contacts, which were related with unwinding events in the protein’s helices. The protein/C_3_N nanosheet interacting process was dominated by vdW contributions to the energy and the stepwise changes observed in the values of this energy term suggested a gradual unfolding pattern of HP35 during the absorption event. Furthermore, we also found that the interaction energy showed a linear correlation with the native Q ratio of HP35, suggesting that the degree of HP35 unfolding was linearly time-dependent to the interaction energy. Our findings shed light on the underlying molecular mechanism of the potential consequences of C_3_N-based nanostructures to proteins, which might delineate the future applications of these nanomaterials in biomedicine.

## INTRODUCTION

The rapid development of a wide variety of nanomaterials have propelled their applications in many scientific areas, including energy conversion, generation of electronic and optical devices, and medicine.^1-9^ Among these and since their discovery,^10-12^ carbon based nanomaterials (CBNs) have received significant attention based on their remarkable mechanical, optical, and electrical properties^4, 13-14^, and particularly in the biomedical field,^15-16^ where they have been applied as possible drug and gene delivery carriers,^17^ in optical imaging devices,^18^ and in different medical treatments (nanotherapeutics).^19-21^ Nonetheless, special attention must be considered during the incorporation of such nanostructures into biological systems, due to potential deleterious effects of these exogenous materials on living cells. When integrated with living organisms, the nanomaterials could either disrupt different cellular components (*e.g.*, cell membrane, proteins, and nucleic acids) and signaling pathways or display little effect in the normal function of the cell and living tissues (*i.e.*, biocompatibility). Regardless of the particular outcome, cytotoxic or biocompatible, the exogenous nanomaterials will encounter a variety of biochemical milieus and consequently, different cellular entities which strengthened the significance to investigate the specifics of the nanomaterial-biomolecule interactions. For example, direct interaction of CBN with proteins or nucleic acids will generate toxicity via the denaturation of the biomolecular native structures (*e.g.*, loss of secondary and/or tertiary structures).^22-23^ Also, by occupying the active site of a receptor and hence inhibiting the native donor-receptor interaction, carbon nanotubes (CNT) can interrupt the normal signal transmission via a non-unfolding pathway.^24^ Moreover, graphene nanosheets can penetrate into and extract large amounts of phospholipids from cell membranes due to the strong dispersion interactions resulting in a significant toxicity by these nanostructures—this graphene’s property suggested the use of this nanomaterial as a possible antibacterial agent.^25^ Lastly, due to their strong surface attraction and influence on the structure and function of biomolecules, CBNs often had to be functionalized prior to their utilization in biomedical applications to enhance biocompatibility—usually by surface modifications with polyethylene glycol or serum proteins.^26-27^

Beyond CBNs, transition-metal dichalcogenides (TMD) nanostructures, particularly those based on molybdenum disulfide (MoS_2_),^28^ have recently emerged as another promising nanomaterial. Interestingly, MoS_2_-based nanomaterials shared similar physicochemical characteristics with CBNs, and thus, replication of the success application of CBN in the biomedical field by MoS_2_-based nanostructures was anticipated. Recent studies have demonstrated the application of MoS_2_-based nanomaterials as antibacterial and antifungal agents,^29^ biosensors (based on their unique direct band gap),^30-31^ photothermal and chemotherapeutic agents (high near-infrared absorbance and extensive specific surface area),^32-33^ as well as contrast agents in X-ray tomography imaging.^33^ Given the extensive usages of MoS_2_ nanostructures in biomedicine, our previous investigations exposed the direct interaction of MoS_2_ and biomolecules and found that pristine MoS_2_ had a non-negligible impact on the protein structure due to its strong attraction,^34-37^ implying the potential toxicity of MoS_2_-based nanomaterials.

More recently, the successful synthesis of carbon nitride compounds, such as C_2_N,^38^ g-C_3_N_4_ (graphitic carbon nitride that includes those materials based on heptazine and triazine units), polytriazine imide (PTI),^39^ as well as polyaniline (C_3_N),^40-41^ provided a novel class of nanomaterials for potential biomedical applications. In this regard, the ultrathin g-C_3_N_4_ nanosheet prepared by a “green” liquid exfoliation route, has been applied as bioimaging candidate with various advantages, including, enhanced intrinsic photoabsorption and photoresponse, high stability, good biocompatibility, and nontoxicity.^42^ Moreover, a g-C_3_N_4_-sensitized TiO_2_ nanotube layer system was produced as visible-light activated efficient metal-free antimicrobial platform.^43^ In addition, the folic acid modified C_3_N dots were selectively endocytosed by and killed tumor cells.^44^ Similarly, 2D C_3_N could be utilized as biosensor.^45^ Despite their diverse applications in biological systems, so far there are very limited studies or evidences to disclose the interaction dynamics and/or potential molecular mechanism of biomolecules binding to carbon nitride compounds. Our recent work probed the adhesion of the prototypical protein villin headpiece (HP35) to a C_2_N nanosheet and found that the protein’s structural integrity was well retained with weaken transverse migration on the C_2_N surface.^46^ Herein, we extended the molecular characterization of carbon nitride compounds to include C_3_N(polyaniline)-based nanostructures. By employing a 2-dimensional C_3_N nanosheet and HP35, we explored the structural consequences at the nano-bio interface of this system—detailed knowledge of the interaction dynamics and molecular mechanism, which was fundamental for its potential applications in biomedicine. Our results revealed that the HP35 protein partially unfolded once absorbed onto the C_3_N nanosheet, which suggested that this carbon-nitrogen-based nanomaterial might have potential influence in modifying biomolecular structures and consequently, potential toxicity to living tissues.

## RESULTS

The 2-dimensional graphene-like C_3_N (also named as polyaniline) nanosheet consisted of the basic unit formed by a benzene ring surrounded by six nitrogen atoms, which was polymerized by anilines^40-41^ (as shown in **Fig. 1a**). HP35, a 35-residue subdomain of the villin headpiece (as shown in **Fig. 1b**), was a small ultrafast folding protein that has been extensively studied by experiments, theory, and simulations.^47-48^ Herein, we employed the well-studied HP35 protein as model system to explore the possible consequences of the presence of the C_3_N nanosheet to biomolecule. Two systems (a typical system configuration was displayed in **Fig. 1c**) were investigated by three independent 1000-ns-length atomistic MD simulations.

**Figure 1.**
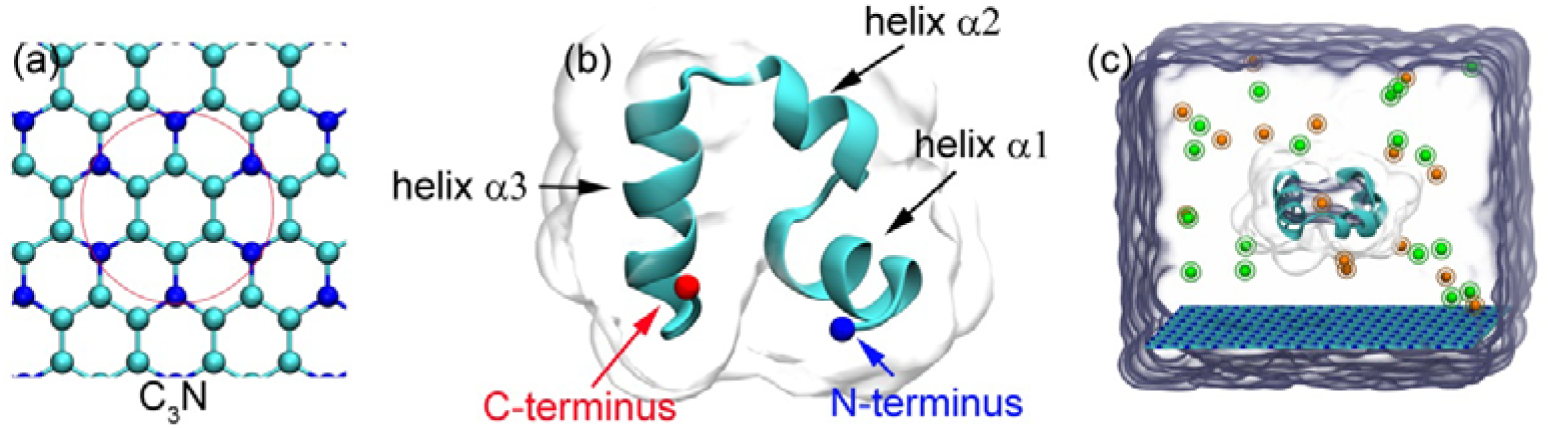
Configurations and simulation setup. (a) Structural conformation of C_3_N. Atoms in the red circle were the elementary unit of C_3_N (i.e., polyaniline). Carbon and nitrogen atoms were represented as cyan and blue spheres, respectively. (b) Ribbon representation of the structure of HP35. The N- and C-termini were indicated by blue and red balls. The three helices composing the protein structure were also labeled as helix α1, helix α2 and helix α3 from the N- to the C-terminus. (c) Representation of the simulated systems investigated in this study. The HP35 protein was placed above the C_3_N nanosheet and rotated by 180° to form two different initial protein orientations. Their initial minimum distance was set to 1.5 nm. Na^+^ and Cl^-^ were depicted as orange and green spheres. The C_3_N sheet was shown with sticks. The water surface showed at the boundaries of the simulated periodic cell.

**Figure 2a** illustrated the relation between the protein/C_3_N nanosheet interaction energy and the changes in the native protein structure. Notably, stronger interactions between the HP35 and C_3_N were directly related with larger structural alterations of the protein structure. Moreover, the loss of the HP35 helical content of its secondary elements (helices α1, α2, and α3) occurred in two of the cases, with remaining helical content values of 40.2±13.9 % and 61.5±3.8 %. The representative structures of HP35 in the insets of **Fig. 2a**, exemplified conformations where the complete loss of helix α1 and helix α2 was observed. The contact probabilities (**Fig. 2b**) of each residue bound to C_3_N along the simulations showed that the residues L1, S2, D3, E4, S15, N19, L27, and F35, were particularly important in mediating the absorption of HP35 onto the surface. Note that the F35 residue at the C-terminus, presented the largest contact probability, up to 78.9%. Detailed analysis demonstrated that F35 formed a face-to-face pattern with the C_3_N surface, yielding a direct interaction energy of −16.4±1.0 kcal/mol (**Fig. 2c**), which suggested its relevance in the binding process—equivalent interactions were observed in the case of HP35 and graphene.^22^

**Figure 2.**
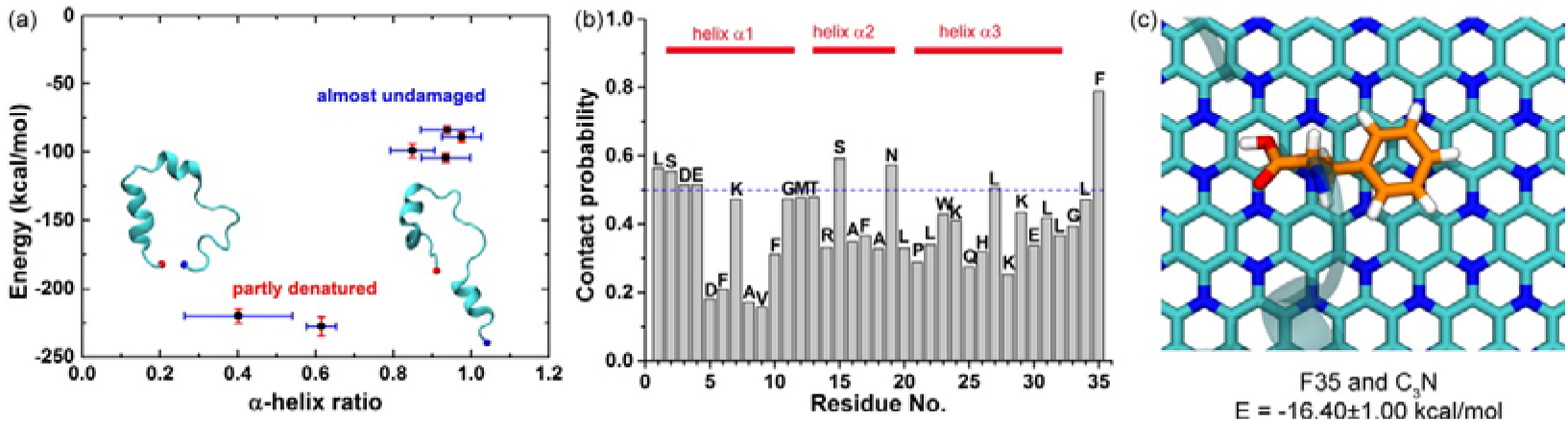
Energy/structure relation and residue contribution. (a) The relation between the interaction energy (including vdW and Coulomb energies between protein and C_3_N) and the secondary structural changes of HP35 collected from the last 50 ns of each trajectory. The inserted HP35 configurations were extracted from two “partly denatured” simulations (see main text). (b) Contact probability for each residue to the C_3_N surface computed by all trajectories. A residue contact was considered if any heavy atom (non-hydrogen atom) of one residue had a distance to the C_3_N surface smaller than 6.0 Å. The F35 residue exhibited the largest contact value, which was an indication of its central role in the absorption event. (c) Illustration of the interactions observed for F35 onto the C_3_N sheet selected from a representative trajectory. The mean interaction energy between F35 and C_3_N was calculated from the last 50 ns.

Next, we performed a dihedral principal components analysis (dPCA) of HP35 bound to C_3_N (**Fig. 3**). Notably, from 7 binding protein conformations on the C_3_N surface extracted from the dPCA plot (namely, conformations i to vii), in 5 of them, HP35 seemingly maintained its structural integrity. However, in two conformations, partial denaturation was observed (conformation ii and iii), that is, the secondary structure of helix α2 was lost in conformation ii while the secondary structure of helix α1 was lost in conformation iii (also see **Fig. 2a**). This finding evidenced the potential denaturation capacity of the C_3_N nanostructures to the HP35 protein.

**Figure 3.**
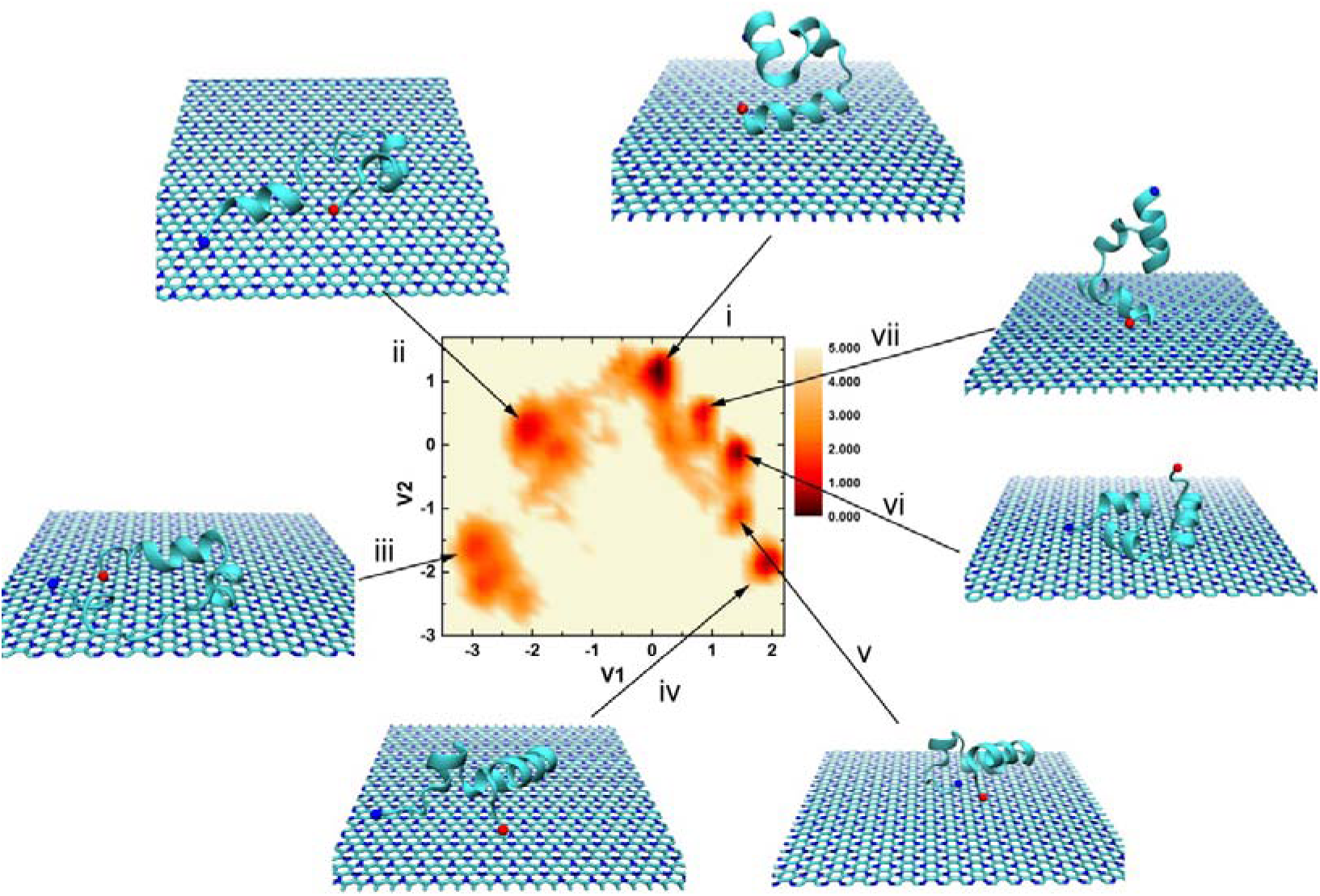
Dihedral principal components analysis (dPCA) of HP35. Projection of the free energy surface for the interaction of HP35 with the C_3_N nanosheet in terms of the two lowest eigenvectors from the dPCA, obtained from the 6 independent 1000 ns trajectories. The color scale bar was in unit of kcal/mol. The key snapshots (including the HP35 and C_3_N nanosheet) were plotted based on the free energy basins, which were labeled in the pictures.

Then, detailed analyses were further conducted to quantitatively measure structural changes of HP35 in binding to the C_3_N surface by selecting a representative trajectory (**Fig. 4**). A root-mean-square deviation (RMSD) calculation of the HP35 protein reflected a substantial increment from its initial value during the 1000 ns simulation time, revealing the significant influence of the C_3_N surface (**Fig. 4a**). Moreover, the number of internal hydrogen bonds in HP35 declined from ∼30 to ∼14 and the native Q value was reduced to 57% (**Fig. 4c**). The time-dependent Q change of each residue showed that most residues lose their native contacts with nearby residues (**Fig. 4c**), particularly the residues V9, G11, and F35 (with ΔQ = 1.0). Furthermore, a map with the time-evolution secondary structure content illustrated the entire loss of helix α1 and the partial loss of helix α3 (**Fig. 4d**). Combined, the quantitative calculations further confirmed the significant depletion of the structural integrity of HP35 upon binding to the C_3_N nanosheet.

**Figure 4.**
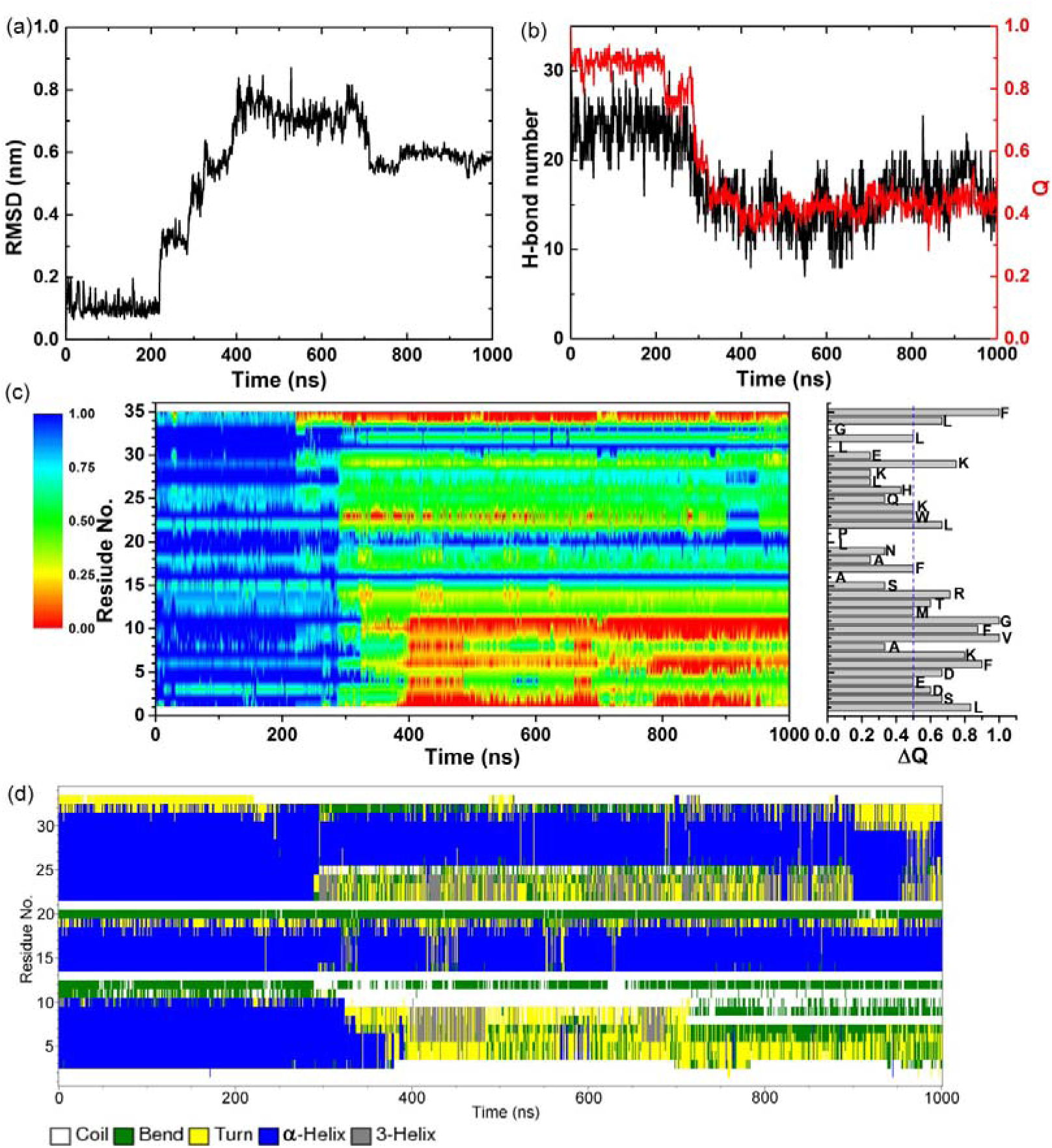
Structural change analyses from a typical trajectory. (a) Root mean square deviation (RMSD) values of HP35 with respect to the simulation time. (b) Time-evolution hydrogen bond number and native contact Q were depicted in black and red, respectively. (c) Time-dependent Q values for each HP35 residue during the simulation (left) and the Q difference value between 0 ns and 1000 ns for each residue (right). (d) The secondary structure map *versus* the simulation time.

To trace the binding kinetics of the system, the relevant aforementioned conformations were depicted based on the change in contact number and in interaction energy (**Fig. 5**). Both, the contact number value and vdW interaction energy exhibited stepwise changes, implying that the HP35 discretely and gradually unfolded and bound to the C_3_N nanosheet, respectively. More importantly, the vdW contributions dominated the interaction throughout the adsorption process. Mechanistically, at t = 13 ns, the HP35 established direct contact via the helix α1 as the contact number value rose to ∼43 and the vdW energy declined to −53 kcal/mol. After 212 ns simulation (t = 225 ns), the orientation of HP35 on the surface slightly changed via a rotation, which bring the F35 residue at the C-terminus in contact with the C_3_N nanosheet. This last event was accompanied by an increment in the contact number value, ∼63, and a decrease in the vdW energy, −82.4 kcal/mol. At this time point, the native structure of HP35 was well maintained without any obvious structural change. However shortly after, at t = 291 ns, the HP35 lay on the C_3_N surface with its three helices all contacting to C_3_N nanosheet in a parallel way with the concomitant loss of the HP35 native structure (exposure and disruption of its hydrophobic core). Meanwhile, the aromatic tryptophan residue, W23, in helix α3 was also positioned in proximity of the C_3_N surface. Lastly, after t = 400 ns, we observed a significant loss in the protein helical content attributed to the complete and partial unwinding of helix α1 and helix α3, respectively. Additionally, direct interaction of several important residues on helix α1—L1, K7, A8 and F10—with the C_3_N nanosheet was established and maintained for the rest of the simulation. Remarkably, we observed a central role of aromatic and basic residues in the surface-binding event, which was also identified in the protein/graphene interaction.^49-50^

**Figure 5.**
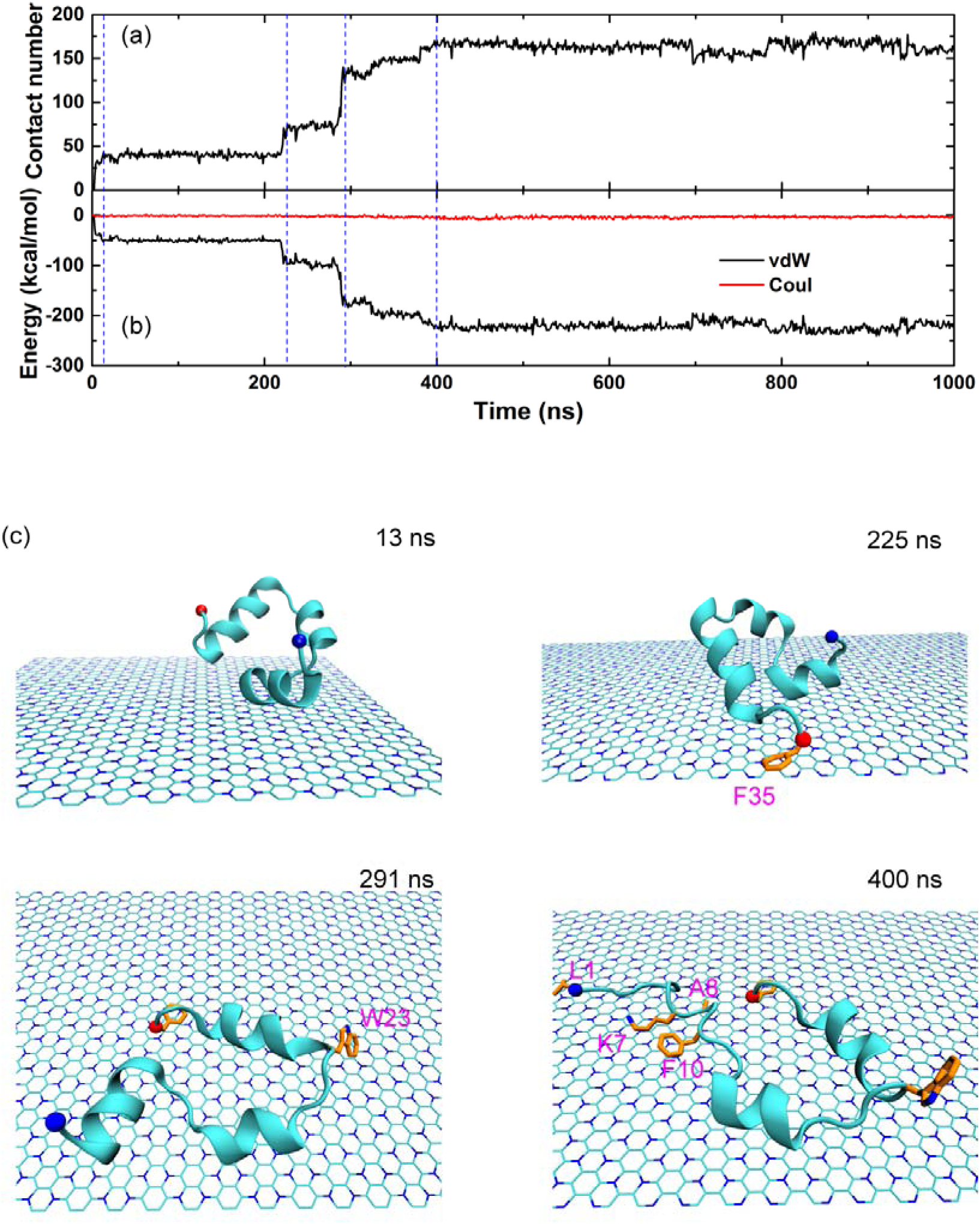
Kinetics of HP35 unfolding on C_3_N nanosheet. Time-evolution contact number (a) and interaction energies (including vdW and Coulomb contributions) of the HP35 absorption process to the C_3_N nanosheet. Blue dashed lines indicated significant time points for the adsorption and unfolding events of HP35. (c) Key snapshots depicted the binding and unfolding processes. Some important residues described in the main text were also displayed.

Finally, we probed the correlation between the interaction energy and native Q ratio in the two representative trajectories where the HP35 unfolding processes were observed, conformation ii and iii (**Fig. 6** and **S1**). Notably, the interaction energy and native Q ratio displayed a time-dependent linear correlation, implying that the unfolding of HP35 on the C_3_N surface had a highly linear correlation with respect to the interaction energy.

**Figure 6.**
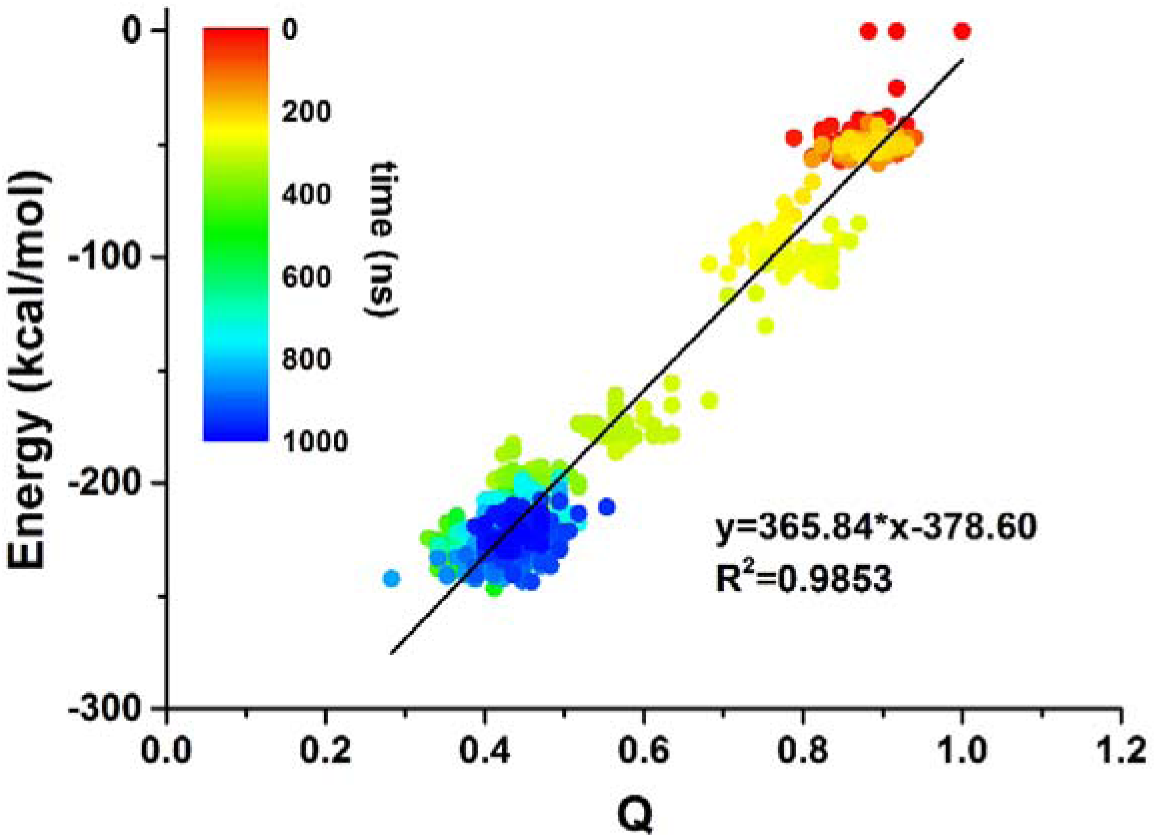
Time-dependent linear correlation of interaction energy and native Q ratio. The color dots were collected every nanosecond (i.e., 1000 dots).

## CONCLUSION

In summary, we explored the potential influence of a C_3_N nanosheet into the structure of the HP35 protein. Our results showed that HP35 was largely denatured upon adsorption to the C_3_N nanosheet. Statistical contact probability demonstrated that the C-terminal F35 residue was a central residue during the surface-binding event. dPCA suggested that in two out of the seven conformations explored by HP35, a significant reduction in the structural integrity of the protein occurred. Further structural analyses quantitatively described the loss in the number of interior hydrogen bonds and native contacts. The HP35 bound to C_3_N surface in a stepwise fashion, which was driven by the vdW energy dominant contribution. The aromatic and basic residues exhibited a critical role during the adsorption and unfolding processes. Moreover, the interaction energy and native Q ratio displayed a time-dependent linear correlation. Our results confirmed the structural influence of the C_3_N nanosheet to protein, which thereby implied a potential toxicity of C_3_N nanomaterial and the need to functionalize its surface to increase biocompatibility for biomedical application.

## METHODS

The C_3_N nanosheet monolayer, treated as benzene rings surrounded and connected by six nitrogen atoms, was displayed in **Fig. 1a**. The C_3_N model utilized in this study consisted of 1,152 carbon and 384 nitrogen atoms with a surface dimension of 5.83×6.73 nm^2^. The 35-residue chicken villin headpiece subdomain protein (HP35, PDB code: 1YRF)^51^ was selected because of its general properties associated with common globular proteins despite its small size.^52-55^ The native structure of HP35 contained a globular bundle of three α-helices, namely helix α1, helix α2 and helix α3, from N-terminus to C-terminus, as shown in **Fig. 1b**. The HP35 was originally placed above the C_3_N nanosheet with their initial distance at 1.5 nm. In this study, we built two systems (one representation was illustrated in **Fig. 1c**) in which the HP35 was rotated for 180° yielding two different orientations relative to the C_3_N surface. The system boxes were set to 5.83×6.73×4.92 nm^3^ and 5.83×6.73×4.90 nm^3^ containing 5,729 and 5,732 water molecules, respectively. In addition, 0.15 M NaCl were also dissolved in two systems to mimic physiological environment.

The simulations were carried out with the GROMACS software package (version 4.6.6)^56^ using the CHARMM36 force field.^57^ The force field for C_3_N was obtained from a recent work.^58^ The VMD software^59^ was used to analyze and visualize the simulation results. The TIP3P water model^60^ was adopted to treat water molecules. Following similar protocols in our previous studies^61-67^, the temperature were maintained at 300 K using v-rescale thermostat^68^ and pressure was kept at 1 atm using semiisotropic Berendsen barostat^69^ (only applied at Z direction perpendicular to the C_3_N nanosheet). To avoid the “artificial collapsing” of nanosheets with their mirror images due to the limited size of simulation box (which was due to the limited computational resources), the C_3_N nanosheet was fixed throughout the simulation process. The long-range electrostatic interactions were treated with PME method,^70^ and the van der Waals (vdW) interactions were calculated with a cutoff distance of 1.2 nm. All solute bonds associated with hydrogen atoms were maintained constant at their equilibrium values with the LINCS algorithm,^71^ and water geometry was also constrained using the SETTLE algorithm.^72^ During the production runs, a time step of 2.0 fs was used, and data were collected every 50 ps. Each system ran for three 1000 ns trajectories. The total aggregated simulation time was about 6 μs.

## ACKNOWLEDGEMENT

We thank Zhi He and Jianxiang Huang for help with the manuscript. This work is partially supported by the National Natural Science Foundation of China (Grants 11574224 and U1967217) and China Postdoctoral Science Foundation (Grant 2019M652069 and 2019T120506). R.Z. also acknowledges the financial support from W. M. Keck Foundation (Grant award 2019-2022).

