## supplementary figure S1 for "Structural Consequences of the Villin Headpiece Interaction with a Carbon Nitride Polyaniline (C_3_N) Nanosheet"

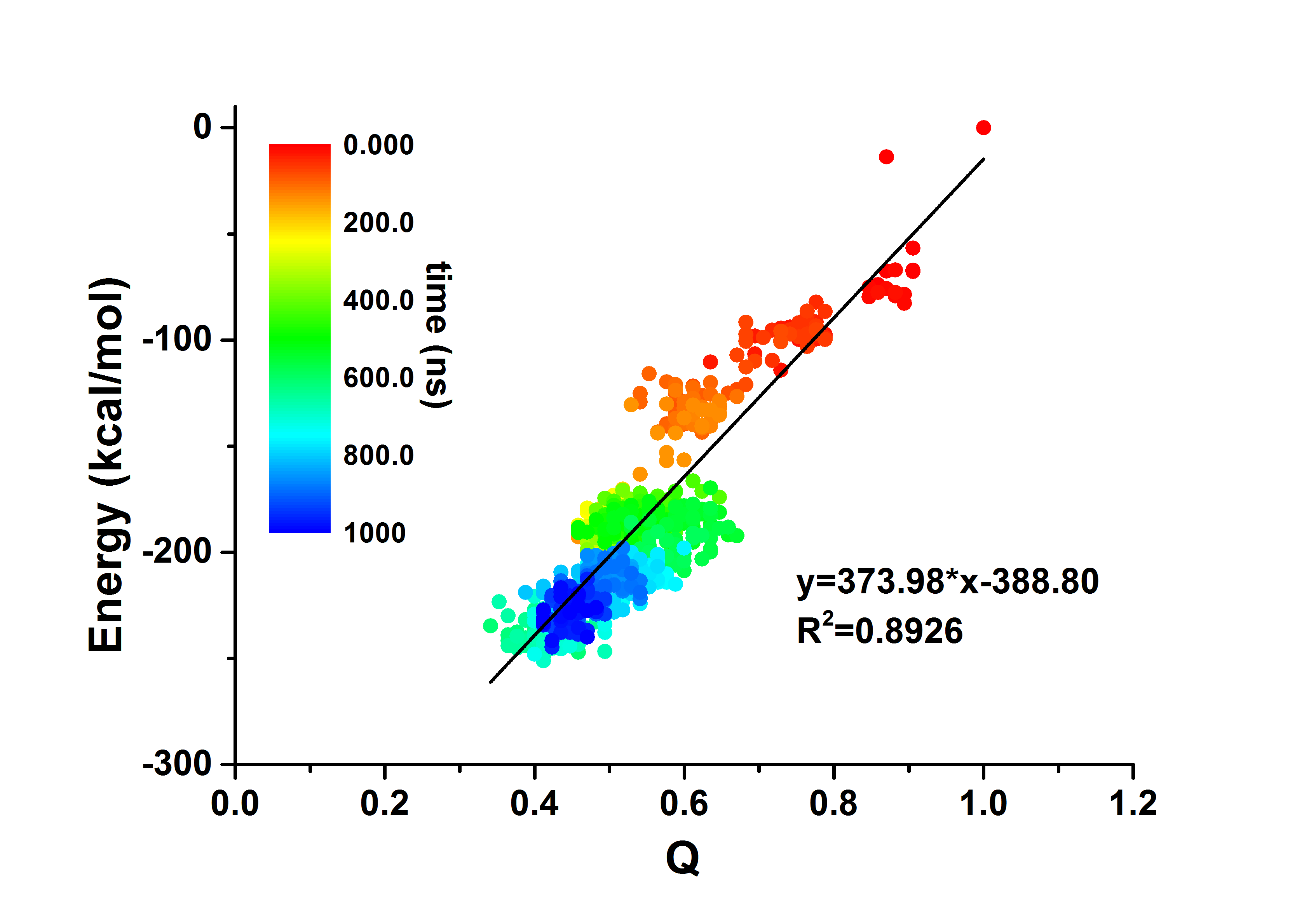

Figure S1. Linear relation between the interaction energy and the structure-dependent Q ratio from a different trajectory as the one in Figure 6 in the main text.
